# Distinctive Gene and Protein Characteristics of Extremely Piezophilic *Colwellia*

**DOI:** 10.1101/2020.03.15.992594

**Authors:** Logan M. Peoples, Than S. Kyaw, Juan A. Ugalde, Kelli K. Mullane, Roger A. Chastain, A. Aristides Yayanos, Masataka Kusube, Barbara A. Methé, Douglas H. Bartlett

## Abstract

**Background:** The deep ocean is characterized by low temperatures, high hydrostatic pressures, and low concentrations of organic matter. While these conditions likely select for distinct genomic characteristics within prokaryotes, the attributes facilitating adaptation to the deep ocean are relatively unexplored. In this study, we compared the genomes of seven strains within the genus *Colwellia*, including some of the most piezophilic microbes known, to identify genomic features that enable life in the deep sea.

**Results:** Significant differences were found to exist between piezophilic and non-piezophilic strains of *Colwellia*. Piezophilic *Colwellia* have a more basic and hydrophobic proteome. The piezophilic abyssal and hadal isolates have more genes involved in replication/recombination/repair, cell wall/membrane biogenesis, and cell motility. The characteristics of respiration, pilus generation, and membrane fluidity adjustment vary between the strains, with operons for a *nuo* dehydrogenase and a *tad* pilus only present in the piezophiles. In contrast, the piezosensitive members are unique in having the capacity for dissimilatory nitrite and TMAO reduction. A number of genes exist only within deep-sea adapted species, such as those encoding d-alanine-d-alanine ligase for peptidoglycan formation, alanine dehydrogenase for NADH/NAD^+^ homeostasis, and archaeal methyltransferase for tRNA modification. Many of these piezophile-specific genes are in variable regions of the genome near genomic islands, transposases, and toxin-antitoxin systems.

**Conclusions:** We identified a number of adaptations that may facilitate deep-sea radiation in members of the genus *Colwellia,* as well as in other piezophilic bacteria. An enrichment in more basic and hydrophobic amino acids could help piezophiles stabilize and limit water intrusion into proteins as a result of high pressure. Variations in genes associated with the membrane, including those involved in unsaturated fatty acid production and respiration, indicate that membrane-based adaptations are critical for coping with high pressure. The presence of many piezophile-specific genes near genomic islands highlights that adaptation to the deep ocean may be facilitated by horizontal gene transfer through transposases or other mobile elements. Some of these genes are amenable to further study in genetically tractable piezophilic and piezotolerant deep-sea microorganisms.

## Background

The deep biosphere makes up one of the largest biomes on earth. An inherent environmental parameter present throughout deep-oceanic and subsurface habitats is high hydrostatic pressure. Elevated hydrostatic pressure influences many aspects of biochemistry and requires adaptations throughout the cell (e.g. Somero, 1992). One well-studied adaptation is the incorporation of unsaturated fatty acids into the membrane to combat physical changes such as decreased fluidity (e.g. DeLong & Yayanos, 1985; DeLong & Yayanos, 1986; Allen *et al*., 1999). Additional membrane-associated adaptations are linked to porin-mediated nutrient transport (Bartlett *et al*., 1989; Bartlett & Chi 1994), respiration (e.g. Yamada *et al*., 2000; Vezzi *et al*., 2005; Xiong *et al*., 2016), and flagellar function (Eloe *et al*., 2008). Within the cell changes in DNA replication, DNA structure, protein synthesis, and compatible solutes are also important (El-Hajj *et al*., 2009; Martin *et al*., 2002; Lauro *et al*., 2008; Yancey *et al*., 2014).

Pressure-induced changes in transcription implicate additional functions (e.g. Campanaro *et al*., 2012; Michoud & Jebbar, 2016). Despite the fact that pressure exerts a profound influence on the nature of life at depth, it is largely ignored in studies of deep-ocean biomes, and in marked contrast to microbial adaptation to temperature or salinity, a robust description of biochemical adaptation to high pressure is lacking.

Only a modest number of psychrophilic (cold-loving) and piezophilic (high-pressure loving) species have been isolated to date, in large part due to the constraints imposed by culturing under under *in situ*, high hydrostatic pressure conditions. However, metagenomic sequencing of deep-ocean communities, and additional analyses of individual microbial genomes, have provided insights. Metagenomic investigations have included locations within the North Pacific subtropical gyre, the Mediterranean and the Puerto Rico Trench (DeLong *et al*., 2006; Martin-Cuadrado *et al*., 2007; Konstantinidis *et al*., 2009; Eloe *et al*., 2011; Smedile *et al*., 2013). Genomic studies include those on *Pseudoalteromonas* (Qin *et al*., 2011), *Alteromonas* (Ivars-Martinez *et al*., 2008), *Shewanella* (Wang *et al*., 2008; Aono *et al*., 2010), *Photobacterium* (Campanaro *et al*., 2005; Vezzi *et al*., 2005; Lauro *et al*., 2014), SAR11 (Thrash *et al*., 2014), and members of the *Thaumarchaeota* (Luo *et al*., 2014; Swan *et al*., 2014). One picture that has emerged from the examinations at this level is that deep-sea microbes are enriched in mobile elements, such as phage and transposases (DeLong *et al*., 2006; Ivars-Martinez *et al*., 2008; Eloe *et al*., 2011; Qin *et al*., 2011; Lauro *et al*., 2013; Smedile *et al*., 2013; Léon-Zayas *et al*., 2015). This has been attributed to the relaxation of purifying selection as an adaptive mechanism (Konstantinidis et al., 2009), either to deep-ocean conditions or to the conditions found on particles (Ganesh *et al*., 2014). Additional properties include an enrichment in heavy metal resistance genes (Ivars-Martinez *et al*., 2008; Eloe *et al*., 2011; Qin *et al*., 2011; Smedile *et al*., 2013; Fontanez *et al*., 2015), the ability to use persistent dissolved organic material under oligotrophic conditions (e.g. Martin-Cuadrado *et al*., 2007; Ivars-Martinez *et al*., 2008; Arrieta *et al*., 2015; Landry *et al*., 2017), and widespread ability for chemoautotrophy (Swan *et al*., 2011; Swan *et al*., 2014; Dyksma *et al*., 2016; Mußmann *et al*., 2017; Pachiadaki *et al*., 2017). The small number of genome sequences of experimentally-confirmed deep-ocean piezophiles include hyperthermophilic archaea (*Pyrococcus* and *Thermoccus*; Vannier *et al*., 2011; Jun *et al*., 2015; Dalmasso *et al*., 2016), a thermophilic bacterium (*Marinitoga*; Lucas *et al*., 2012), a mesophilic bacterium (*Desulfovibrio*; Pradel *et al*., 2013) and psychrophilic bacteria (*Photobacterium*, *Psychromonas,* and *Shewanella*; Vezzi *et al*., 2005; Aono *et al*., 2010; Lauro *et al*., 2013a; Lauro *et al*., 2013b; Zhang et al., 2019b). The genomic adaptations of these microorganisms to the deep ocean or high hydrostatic pressure have not been fully explored (e.g. reviewed in Simonato *et al*., 2006; Lauro *et al*., 2008; Oger & Jebbar, 2010; Peoples & Bartlett, 2017). Thus far the genome characteristics of only one experimentally-confirmed obligately psychropiezophilic bacterial species, *Shewanella benthica* (Lauro *et al*., 2013a; Zhang *et al*., 2019b), and one species of obligately thermopiezophilic archaeon, *Pyrococcus yayanosii* (Jun *et al*., 2011), have been described.

Most known psychropiezophilic strains belong to phylogenetically narrow lineages of *Gammaproteobacteria*, including members of the *Colwellia*, *Shewanella*, *Moritella*, *Photobacterium,* and *Psychromonas* (reviewed in Jebbar *et al*., 2015; Nogi *et al*., 2017). The genus *Colwellia* contains some of the most psychrophilic and piezophilic species currently known. Members of this genus are heterotrophic and facultatively anaerobic (Bowman *et al*., 2014). This genus has been of recent interest because of its association with the Deepwater Horizon oil spill, where members of the *Colwellia* became some of the most abundant taxa present because of their ability to degrade hydrocarbons (Redmond & Valentine, 2012; Mason *et al*., 2014; Kleindienst *et al*., 2015). Although *Colwellia* do not appear to be abundant members of deep-ocean or hadal (typically < 1%; e.g. Eloe *et al*., 2011; Tarn *et al*., 2016; Peoples *et al*., 2018) communities, they can become dominant members under mesocosm conditions (Hoffmann *et al*., 2017; Boeuf *et al*., 2019; Peoples *et al*., 2019a). At least four piezophiles have been successfully isolated and described from this genus. The first known obligate psychropiezophile, originally designated *Colwellia* sp. MT41, was isolated from the amphipod *Hirondellea gigas* from the Mariana Trench at a depth of 10,476 m (Yayanos *et al*., 1981). Strain MT41 shows optimum growth at 103 MPa and does not grow at a pressure below 35 MPa, approximately the pressure at average ocean depths (Yayanos *et al*., 1981; Yayanos, 1986; DeLong *et al*., 1997). Recently, *Colwellia marinimaniae* MTCD1, the most piezophilic microbe known to date, was isolated from an amphipod from the Mariana Trench (Kusube *et al*., 2017). This strain displays an optimum growth pressure of 120 MPa and a growth range from 80 to 140 MPa, higher than the pressure found at full ocean depth. Based on 16S rRNA gene similarity both strains MT41 and MTCD1 strains were determined to belong to the species *Colwellia marinimaniae* (Kusube *et al*., 2017). Other psychropiezophiles within the genus include *C. hadaliensis* (Deming *et al*., 1988) and *C. piezophila* (Nogi *et al*., 2004), isolated from the Puerto Rico and Japan trenches, respectively. While the growth characteristics and fatty acid profiles of these piezophilic species of *Colwellia* have been reported, other adaptations of these strains for dealing with high hydrostatic pressure and deep-ocean environmental conditions have not been investigated in great detail.

In this study, we compared the genomes of members of the *Colwellia* to identify attributes that confer adaptation to the deep ocean. We report the genome sequences of three obligately piezophilic *Colwellia*; *Colwellia marinimaniae* MT41, *C. marinimaniae* MTCD1, and a new isolate obtained from sediment in the Tonga Trench, *Colwellia* sp. TT2012. We compared these genomes, along with the publicly-available genome of *C. piezophila* ATCC BAA-637 (isolated as strain Y223G; Nogi *et al*., 2004), against three piezosensitive strains of *C. psychrerythraea*. The piezosensitive strains include the most well-studied member of the *Colwellia*, *C. psychrerythraea* 34H, a psychrophile isolated from Arctic ocean sediments (Huston *et al*., 2000) whose adaptations to low temperature have been investigated at multiple levels (e.g. Marx *et al*., 2009; Showalter & Deming, 2018), including with genomics (Methé *et al*., 2005). The two other comparison strains are *C. psychrerythraea* GAB14E and ND2E, obtained from the Great Australian Bight at a depth of 1472 m and the Mediterranean Sea from 495 m, respectively (Figure 1A; Techtmann *et al*., 2016). While the *C. psychrerythraea* strains share 99% identical 16S rRNA sequences, they have very divergent average nucleotide identities (ANI; Techtmann *et al*., 2016). Because low temperatures and high pressures have similar effects on biochemical processes, these three microbes were selected as comparison strains because they all show growth at low temperatures, reducing the impact of temperature as a confounding factor. Through the comparison of these seven strains depth and pressure-associated shifts were identified in protein amino acid distributions and isoelectric points, as well as in gene abundances, including the discovery of piezophile-specific genes.

**Figure 1.**
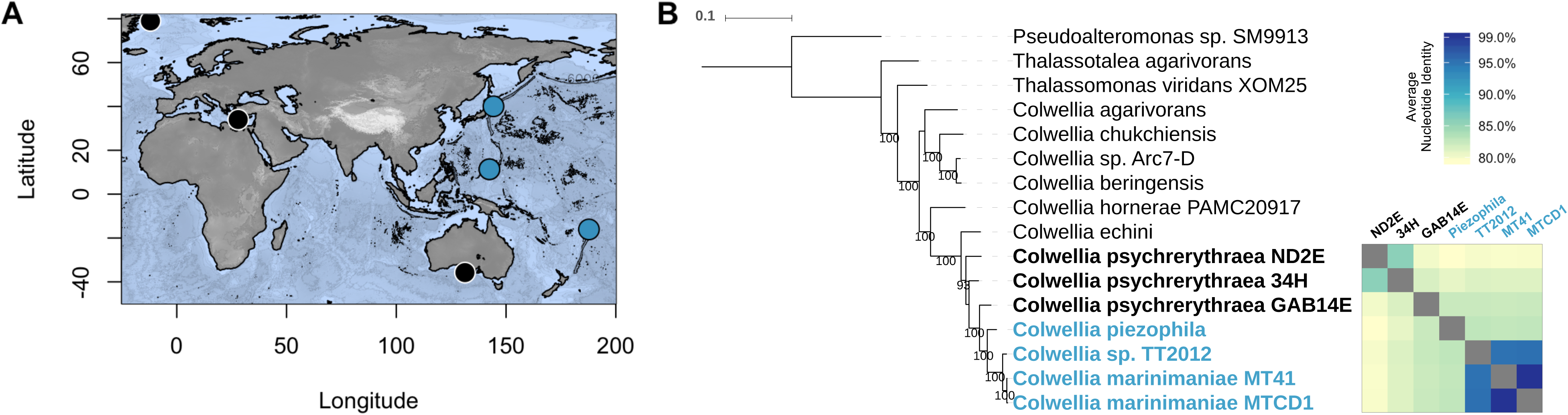
A; Approximate sample collection locations for the *Colwellia* strains compared in this study. B; Whole genome phylogenetic tree and shared average nucleotide identities among the seven strains of interest.

## Results

### General characteristics

We first evaluated the influence of high hydrostatic pressure on the growth of the seven strains of *Colwellia*. The growth characteristics of *Colwellia marinimaniae* MT41, *C. marinimaniae* MTCD1, and *C*. *piezophila* have been previously reported, showing growth optima at 103 MPa (Yayanos, 1986; DeLong *et al*., 1997), 120 MPa (Kusube *et al*., 2017), and 60 MPa (Nogi *et al*., 2004), respectively. *Colwellia* sp. TT2012 is obligately piezophilic, showing growth at 84 and 96 MPa but not at atmospheric pressure. We report the optimum growth pressure in this manuscript as 84 MPa because strain TT2012 was not able to be revived after repeated cryopreservation for growth rate analyses at lower or higher pressures. The three *C. psychrerythraea* strains displayed different growth patterns from one another, but similarly showed no growth at a pressure of 40 MPa after 10 days regardless of temperature (4°C or 16°C; Supplementary Figure 1). Based on these growth characteristics, we classified the microbes as either piezophilic (*C. marinimaniae* MT41, *C. marinimaniae* MTCD1, *Colwellia* sp. TT2012, and *C. piezophila*) or piezosensitive (*C. psychrerythraea* strains 34H, GAB14E, and ND2E). These terms are used to describe these groupings for the remainder of the manuscript.

To identify genomic attributes that facilitate growth at high pressure in the deep sea, we compared the genomes of the piezophilic and piezosensitive strains (Table 1). We report here for the first time the genome sequences of *Colwellia marinimaniae* MT41, *C. marinimaniae* MTCD1, and *Colwellia* sp. TT2012. The remaining genomes are either publicly available (*C. piezophila*, Kyrpides *et al*., 2014) or have been previously reported (strain 34H, Methé *et al*., 2005; strains ND2E and GAB14E, Techtmann *et al*., 2016). The piezophiles are more closely related to one another than to the piezosensitive strains based on a whole genome marker tree and average nucleotide identity (Figure 1). This is also true when the strains are compared using a ribosomal 16S RNA gene phylogenetic tree (Supplementary Figure 2). *Colwellia marinimaniae* MT41, *C. marinimaniae* MTCD1, and *Colwellia* sp. TT2012 share approximately 96% 16S rRNA gene sequence similarity and formed a monophyletic clade with an isolate from the Kermadec Trench. Despite being isolated 34 years apart, strains MT41 and MTCD1 share greater than 99% 16S rRNA gene sequence similarity and ANI. In contrast, the ANI of these strains are only 95% similar to TT2012, indicating that TT2012 represents a distinct species. *C. piezophila* does not appear to belong to this 16S rRNA gene tree piezophile-only monophyletic clade (Supplementary Figure 2), although this relationship could not be confirmed with a whole genome marker tree due to a lack of related genomes. Despite showing greater than 98% 16S rRNA gene sequence similarity, the ANI of *C. psychrerythraea* strains 34H, GAB14E, and ND2E is less than 90%, indicating that they have highly variable genome sequences.

**Table 1.** Genome characteristics of strains of *Colwellia* compared in this study.

| Strain | Isolation location | Isolation depth | Isolation source | Genome size (Mb) | DNA scaffold count | Completeness | Contamination | GC | Coding region | Predicted genes | Protein coding genes with function prediction |
| --- | --- | --- | --- | --- | --- | --- | --- | --- | --- | --- | --- |
| <u>MTCD1</u> | Mariana Trench | 10918 m | Amphipod | 4.37 | 184 | 100% | 1.47% | 39.34% | 83.68% | 3895 | 2826 |
| <u>MT41</u> | Mariana Trench | 10476 m | Amphipod | 4.34 | 1 | 100% | 0.73% | 39.40% | 83.88% | 4057 | 2933 |
| <u>TT2012</u> | Tonga Trench | 9161 m | Sediment | 4.44 | 250 | 99.33% | 2.61% | 39.55% | 83.00% | 4071 | 2897 |
| <u>C. piezophila</u> | Japan Trench | 6278 m | Sediment | 5.48 | 38 | 100% | 1.01% | 38.84% | 83.65% | 4598 | 3362 |
| <u>GAB14E</u> | Great Australian Bight | 1472 m | Water | 5.72 | 77 | 99.49% | 0.68% | 37.97% | 85.76% | 4790 | 3484 |
| <u>ND2E</u> | Mediterranean Sea | 495 m | Water | 5.15 | 57 | 100% | 2.38% | 38.08% | 85.73% | 4479 | 3379 |
| <u>34H</u> | Arctic Ocean | 305 m | Sediment | 5.37 | 1 | 100% | 1.68% | 38.01% | 85.81% | 5066 | 3233 |

### GC content and amino acid features

We first compared general genomic attributes of the piezophilic and piezosensitive strains, including genome size, GC content, isoelectric point, and amino acid distribution. Genome sizes ranged between 4.3 and 5.7 Mbp in size (Table 1). The three piezophiles isolated from the deepest depths (strains MT41, MTCD1, TT2012) have smaller genomes than the piezosensitive strains (T-test, p < .031), but no correlation between genome size and optimum growth pressure was found when considering *C. piezophila* and other members of the *Colwellia* (Supplementary Figure 3). Coding density is lower in the piezophilic *Colwellia*. This is true even when including all sequenced members of the *Colwellia* (T-test, p < .01). GC content ranged between ∼ 38 and 39%, with slightly higher GC present in the piezophiles. This is also true when compared with other *Colwellia* strains with the exception of *C. chukchiensis* (Supplementary Figure 3; T-test, p < .08). However, when examining members of the genera *Colwellia*, *Psychromonas*, and *Shewanella*, no correlation was apparent between % GC and growth pressure. No correlation was found between optimum growth pressure and % GC within full length 16S rRNA genes in the *Colwellia*.

Next, we evaluated the isoelectric point distributions of the *Colwellia*. Both piezophilic and piezosensitive strains show a similar bimodal distribution of protein isoelectric points. However, the piezophiles have a higher number of basic proteins (Figure 2; T-test, p < .01). This shift is also seen when comparing within a broader number of *Colwellia* (T-test, p < .01) with the exception of *C. chukchiensis* (Supplementary Figure 4). Piezophilic strains within the genera *Psychromonas* and *Shewanella* also show a higher number of basic proteins compared to their piezosensitive counterparts (Supplementary Figure 4; T-test, Psychromonas, p < .03; T-test, Shewanella, clade 3, p < .34), with obligate piezophiles such as *Shewanella benthica* KT99, *Psychromonas* sp. CNPT3, and an uncultured *Psychromonas* single-amplified genome from a hadal amphipod (Leon-Zayas *et al*., 2015) having dramatically more basic proteins. GC content or optimum growth temperature does not appear to be responsible for this shift in pI bias, even when taking into account within-genus phylogenetic clades (Supplementary Figure 4, Supplementary Figure 5).

**Figure 2.**
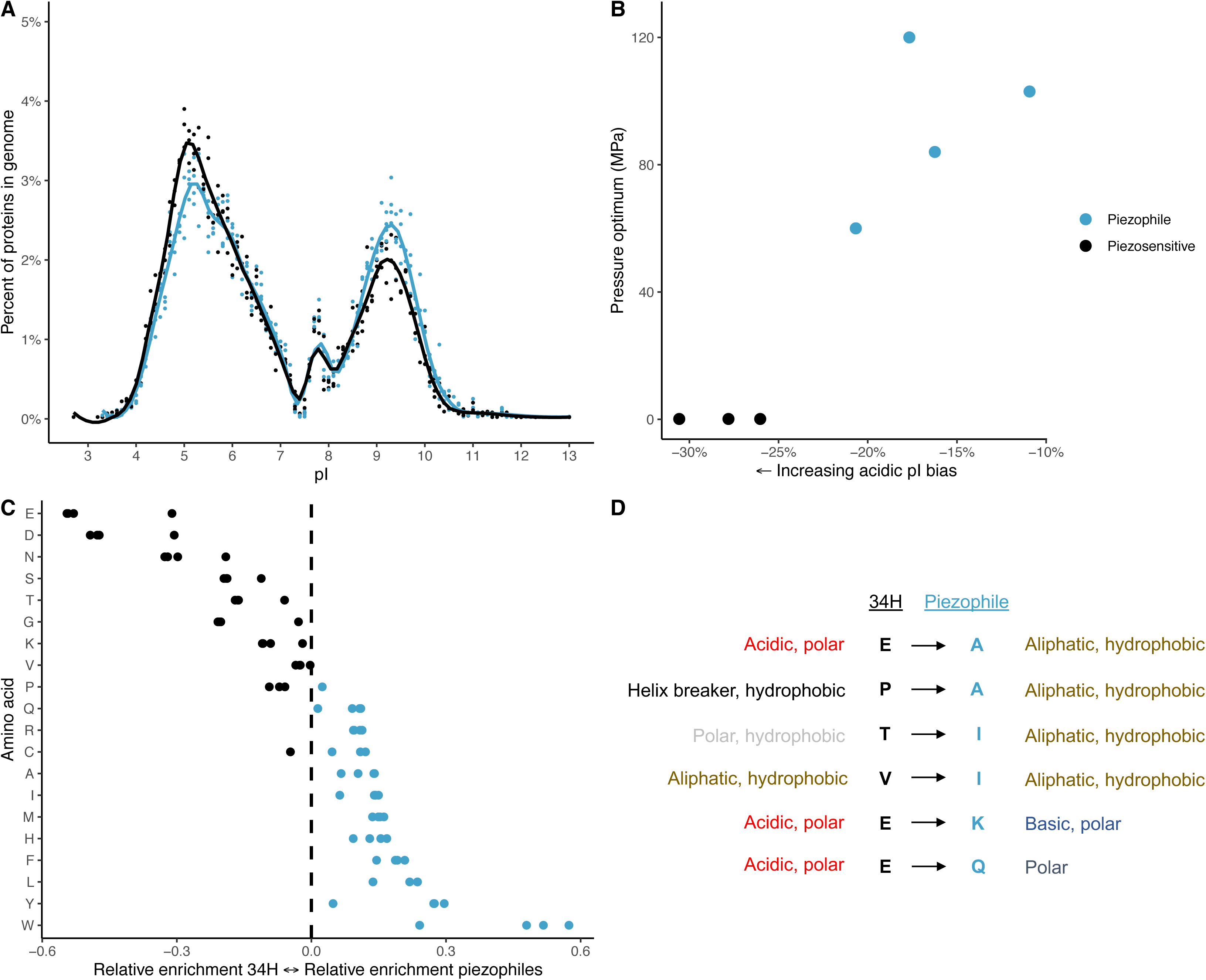
A; Isoelectric point distribution of proteins within piezophilic (blue points) or piezosensitive (black) strains, with an average line of fit within each group. B; Isoelectric point protein bias within each strain as a function of their growth pressure. C; Asymmetry index values indicating preference of amino acids in the piezophiles or *C. psychrerythraea* 34H within orthologous proteins present in all strains. D; Specific amino acid substitutions from *C. psychrerythraea* 34H to the piezophiles within orthologous proteins. The substitutions shown were also identified within comparisons between piezophilic and piezosensitive *Shewanella*.

Comparisons of amino acid abundances within conserved, orthologous proteins showed that certain amino acids are more abundant in the piezophilic proteins when compared to those in *C. psychrerythraea* 34H (Figure 2). Amino acids that are specifically enriched in the piezophiles included tryptophan (W), tyrosine (Y), leucine (L), phenylalanine (F), histidine (H), and methionine (M). In contrast, amino acids enriched in the piezosensitive strain included glutamic acid (E), aspartic acid (D), asparagine (N), and serine (S). Specific amino acid asymmetrical substitutions in which one amino acid consistently replaced another, including substitutions that were also conserved in comparisons within members of the *Shewanella*, from piezosensitive to piezophilic amino acid were: glutamic acid → alanine (A), proline (P) → alanine, threonine (T) → isoleucine (I), valine (V) → isoleucine (I), glutamic acid → lysine (K), asparagine (N) → lysine, glutamic acid → glutamine (Q; Figure 2). Further asymmetrical substitutions specific to the genus *Colwellia* include, from non-piezophile to piezophile, aspartic acid → alanine, glycine (G) → alanine, serine → alanine, asparagine → histidine, valine → leucine, and glutamic acid → threonine.

### Gene differences

We compared the predicted gene complements of the piezophilic and piezosensitive strains. When comparing relative abundances of clusters of orthologous genes (COGs; Figure 3), piezophilic *Colwellia* have a higher percentage of genes for replication/recombination/repair (Category L), cell wall/membrane biogenesis (Category M), cell motility (Category N), extracellular structures (Category W), and translation and ribosomal structure (Category J). The piezosensitive strains have higher percentages of genes for transcription (Category K), secondary metabolite biosynthesis/transport/metabolism (Category Q), and general function prediction (Category R). Transposable elements are notably more abundant in the piezophiles, with the exception of *C. piezophila*, having almost twice as many transposases as their piezosensitive counterparts (Figure 3). Toxin-antitoxin genes are also enriched in the piezophiles, with piezophilic strains containing 24-33 toxin-antitoxin genes while the piezosensitive *Colwellia* have 9-18 copies. We found that strain MT41 and *C. psychrerythraea* 34H have approximately 11 and 9 genomic islands (GIs), respectively, as determined using Island Viewer (Bertelli *et al*., 2017). We do not report the total number of GIs in the other strains because the fragmentation of their genomes likely leads to GI misidentification.

**Figure 3.**
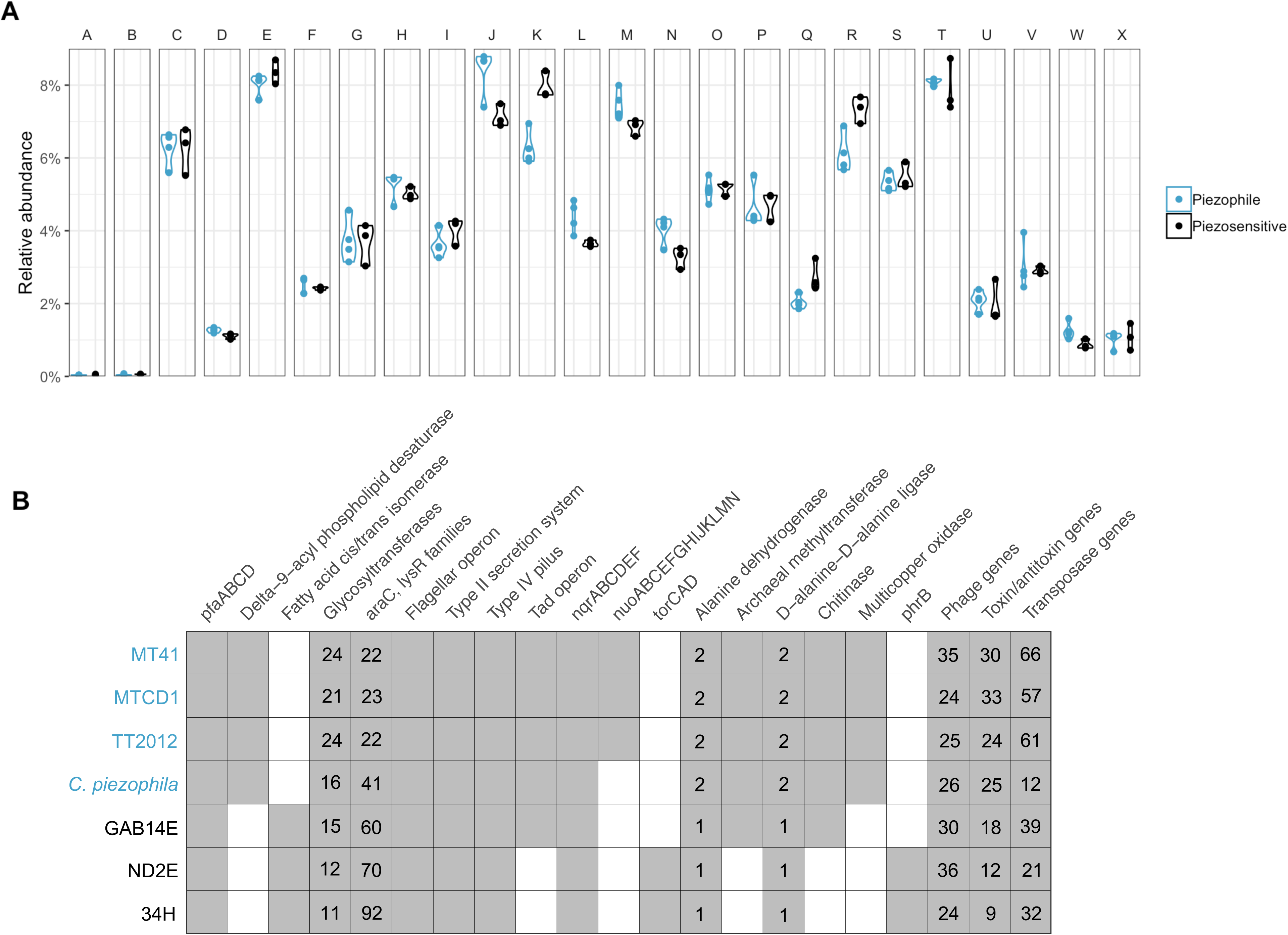
A; The percent abundance of proteins within each COG category within piezophilic or piezosensitive *Colwellia*. B) Specific genomic attributes that were differentially present in piezophilic or piezosensitive strains. Present, grey; absent, white.

All of the strains analyzed are heterotrophic. However, potential differences in carbon metabolism exist (Figure 3). Genes for sarcosine oxidase (*soxBDAG*), which function in the catabolism of glycine betaine in *Colwellia* (Collins & Deming, 2013), are present in 34H and ND2E but not in the piezophiles. Transporters and permeases for putrescine are enriched in 34H and GAB14E, strains where putrescine has been experimentally shown to be used as a sole carbon source (Techtmann *et al*., 2016). In contrast, we identified genes involved in chitin degradation, such as a chitin binding protein and chitinase (family 10 and 18), in the piezophiles and GAB14E but not in the other piezosensitive strains.

Members of the *Colwellia* are facultative anaerobes capable of respiration and fermentation. While all the *Colwellia* compared here use both the rnf (*rnfABCDGE*) and Na^+^-nqr (*nqrABCDEF*) respiratory complexes, the NADH dehydrogenase I complex (*nuoABCEFGHIJKLMN*) is only present in the three hadal piezophiles. These genes show similarity to those in the piezophiles *Shewanella benthica* and *S. violacea* and to metagenomic sequences from hadal sediments (Peoples, 2018). While all seven strains may have the capacity for assimilatory nitrite (such as *nirBD*, *nasA*) and nitrate reduction (*napCBADFE*), genes for dissimilatory nitrite reduction (*nirSCFNTB*) are only present in *C. psychrerythraea* strains 34H and ND2E. The gene *nirK* is present in *C. piezophila*, although this strain was shown to reduce nitrate but not nitrite (Nogi *et al*., 2004). The gene cluster for nitrous oxide reduction, *nosRZDFYL*, is present in strains 34H, ND2E, and *C. piezophila*. This operon is flanked by conserved regions found in the other strains, suggesting an insertion or deletion event. Furthermore, the capacity for trimethylamine-N-oxide reduction using *torSTRECAD* is present in strains 34H and ND2E but not in any of the piezophiles.

The seven strains of *Colwellia* compared are psychrophilic or psychrotolerant and have adaptations to low temperatures. For example, all contain *pfaABCD* to produce polyunsaturated fatty acids to counteract decreases in membrane fluidity because of low temperatures. In the case of the deep-sea *Colwellia* this system will also optimize membrane phospholipid physical state at high pressure. However, a number of genes involved in membrane adaptation are differentially present among the two *Colwellia* groups. All piezophilic *Colwellia* have genes encoding delta-9 acyl-phospholipid desaturase, another enzyme promoting unsaturated fatty acid synthesis by introducing double bonds directly into membrane phospholipid saturated fatty acids. In contrast, a fatty acid cis/trans isomerase that alters the ratio of cis- and trans-phospholipids by isomerizing -cis to -trans double bonds, is encoded within all piezosensitive *Colwellia* but is notably absent in the piezophilic *Colwellia*. Furthermore, the piezophilic strains encode almost twice as many glycosyltranferases, enzymes involved in extracellular polysaccharide synthesis.

Stress-response genes are also differentially present in the genomes. Deoxyribopyrimidine photolyase (DNA photolyase; *phrB*), which is involved in repairing DNA damaged by ultraviolet light, is found in strains 34H and ND2E but notably absent in all piezophilic *Colwellia*. Both piezophilic and piezosensitive strains contain superoxide dismutase and catalase for responding to oxidative stress. The genes *araC* and *lysR*, whose products control the expression of a variety of stress response systems, are more abundant in the piezosensitive *Colwellia*. The piezophilic *Colwellia* are distinct in having multicopper oxidases and copper chaperones for coping with heavy metal damage and maintaining copper homeostasis.

Phenotypic analysis of the *Colwellia* showed that the piezophiles appear more resistant to copper exposure compared to their non-piezophilic counterparts (Supplementary Figure 6). Some of the genes which putatively confer heavy metal resistance are similar to other piezophiles and are located near genomic islands or other horizontally transferred elements, consistent with the hypothesis that heavy metal genes can be horizontally transferred (e.g. Orellana & Jerez, 2011; Navarro *et al*., 2013; Chen *et al*., 2017).

We identified other unique genes that differ not only between *Colwellia* strains but show biased distributions towards additional piezophilic microbes and deep-ocean metagenomic datasets (Table 2; Dombrowski *et al*., 2018; Hu *et al*. 2018; Tully *et al*., 2018; Peoples, 2018). For example, a putative archaeal S-adenosyl-l-methionine (SAM) dependent methyltransferase (pfam13659) is present in the piezophiles and strain GAB14E. This protein is similar to those present in bacterial and archaeal piezophiles, including members of the genera *Colwellia*, *Shewanella*, *Moritella*, *Psychromonas*, *Methanocaldococcus*, *Thermococcus*, and *Pyrococcus*. The related methyltransferase isolated from *Pyrococcus abysii* (39% similar to MT41 protein) functions in tRNA modification (Guelorget *et al*., 2010). Piezophilic *Colwellia* have two copies of d-alanine-d-alanine ligase (pfam07478), a gene which may be involved in peptidoglycan synthesis. Unlike the situation in piezophilic *Shewanella* (Zhang *et al*., 2019b), this gene is not present near flagellar assembly components. While all strains have operons for a Type II secretion system and a Type IV pilus, a *tad* pilus involved in adhesion is found only in the piezophiles and related to that in *Shewanella violacea*. This operon is also found in GAB14E; however, this strain lacks a number of putative tadE-like genes that are present in the piezophile operons. Two alanine dehydrogenases are also present in the piezophilic strains while only one is present in the piezosensitive members. The piezophile-specific dehydrogenase (pfam05222) is thought to catalyze the NAD-dependent reversible amination of pyruvate to alanine. It is similar to a dehydrogenase present in other piezophilic species, including *Shewanella benthica*, *Moritella yayanosii*, *Photobacterium profundum* SS9, and binned genomes from a deep subsea aquifer (Tully *et al*., 2018) and trench sediments (Peoples, 2018).

**Table 2.** Genes identified in piezophilic *Colwellia* but not the piezosensitive strains and which show a biased presence within other known piezophilic microbes and deep-ocean datasets.

| IMG Gene ID MT41 | Start MT41 (bp) | End MT41 (bp) | Similar to: |
| --- | --- | --- | --- |
| 2501712773-2501712774 | 738561 | 741622 | *P. hadalis* , *S. benthica* , *S. violacea* , *M. yayanosii* , *Moritella* sp. PE36, Peoples 2018, Hu *et al.* 2018, Tully *et al.* 2018 |
| 2501712781 | 748798 | 749364 | *S. benthica* , Tully *et al.* 2018, Dombrowski *et al.* 2018 |
| 2501712785 | 751307 | 751420 | *P. hadalis* , Peoples 2018 |
| 2501713024-2501713025 | 1002524 | 1003568 | *M. yayanosii* , *Moritella* sp. PE36, Tully *et al.* 2018, Dombrowski *et al.* 2018 |
| 2501713028-2501713043 | 1004921 | 1020893 | *P. hadalis* , *S. benthica* , *S. violacea* , *M. yayanosii* , *Moritella* sp. PE36, Hu *et al.* 2018, Tully *et al.* 2018 |
| 2501713628 | 1635614 | 1636453 | *P. hadalis* , *S. benthica* , *S. violacea* , *M. yayanosii* , *Moritella* sp. PE36, piezophilic archaea, Hu *et al.* 2018, Tully *et al.* 2018 |
| 2501713976 | 1995082 | 1995321 | *S. benthica* |
| 2501714033 | 2052442 | 2052666 | *S. benthica* |
| 2501714084 | 2101280 | 2101915 | *P. hadalis* , *S. benthica* , *S. violacea* , *M. yayanosii* , *Moritella* sp. PE36, Tully *et al.* 2018 |
| 2501714124-2501714126 | 2137413 | 2141565 | *P. hadalis* , *S. benthica* , *M. yayanosii* , Tully *et al.* 2018, Dombrowski *et al.* 2018 |
| 2501714471-2501714485 | 2514635 | 2530350 | *S. benthica* , *S. violacea* , Peoples 2018 |
| 2501714619 | 2663589 | 2663918 | *S. benthica* , Peoples 2018, Tully *et al.* 2018 |
| 2501714669 | 2714988 | 2715770 | *M. yayanosii* , *Moritella* sp. PE36, SAR324, Peoples 2018, Tully *et al.* 2018 |
| 2501715698 | 3869630 | 3871057 | *Photobacterium profundum* SS9, *S. benthica* , *M. yayanosii* , *Moritella* sp. PE36, Peoples 2018, Tully *et al.* 2018 |
| 2501715722 | 3894109 | 3895707 | *P. hadalis* , Tully *et al.* 2018 |
| 2501715931-2501715932 | 4122279 | 4122819 | *S. benthica* , *M. yayanosii* |
| 2501716002-2501716003 | 4182966 | 4183371 | *S. benthica* , *M. yayanosii* , Tully *et al.* 2018 |

A number of the genes specific to piezophiles are present near one another, rather than individually spread throughout the genome (Table 2). Many of these genes are near variable regions containing genomic islands, phage genes, transposases, and toxin-antitoxin system genes (Supplementary Figure 7). For example, the d-alanine-d-alanine ligase in strain MT41 is next to two putative genomic island regions, one of which is different than that present in strain TT2012 (Figure 4). Because genomic islands are identified based on nucleotide bias across the genome and the *Colwellia* sp. TT2012 genome is fragmented into short contigs, the lack of predicted genomic islands does not preclude their presence. In the piezophile *Moritella yayanosii* this gene is near a gene encoding a predicted phage integrase protein, while in *Shewanella benthica* KT99 it is present in a flagellar operon that also contains a transposase embedded within it. Similarly, the piezophile-specific alanine dehydrogenase is present near a number of phage and toxin/antitoxin genes and downstream from a genomic island. In strain TT2012, this gene is in the middle of a putative genomic island (Figure 5), while in *Photobacterium profundum* SS9 it is flanked on one side by a transposase. Some of the genes present in these variable regions, when not specific to piezophiles, display low similarity to members of the genus *Vibrio*. The similarity of variable genes within *Colwellia* to species of *Vibrio* has been previously noted (Collins & Deming, 2013). Horizontal gene transfer has been shown to be important in the evolution of *Vibrio* species (Faruque & Mekalanos, 2013).

**Figure 4.**
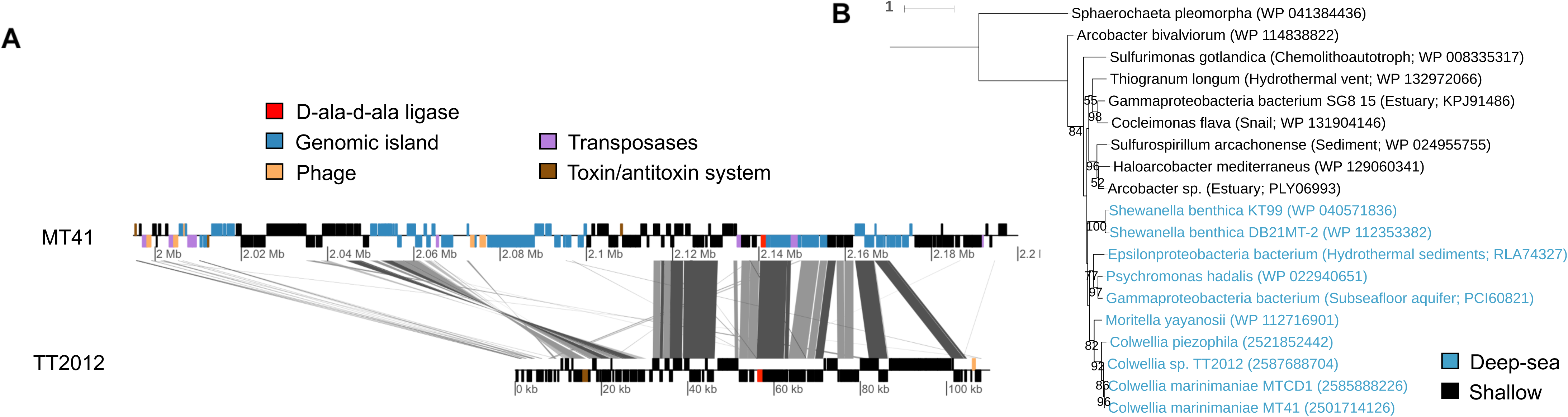
A; The location of a d-ala d-ala ligase gene in strains MT41 and TT2012, with surrounding genes labeled. B; An amino acid tree of the d-ala-d-ala ligase with sequences approximately > 50% similar shown.

**Figure 5.**
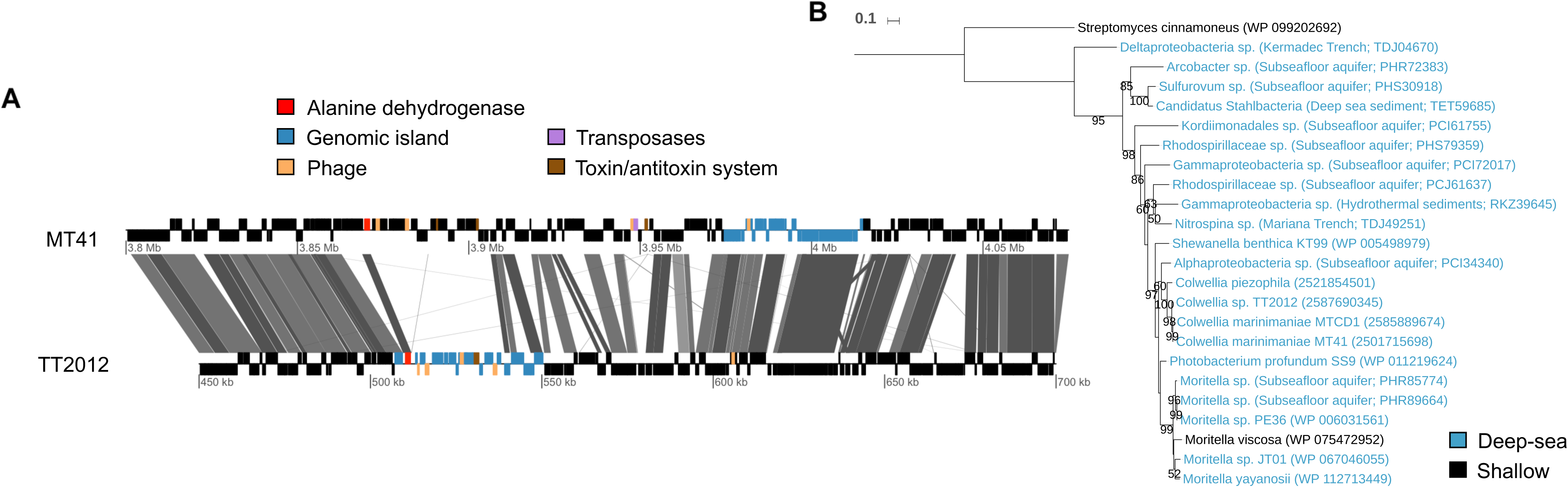
A; The location of alanine dehydrogenase genes in strains MT41 and TT2012, with surrounding genes labeled. B; An amino acid tree of the alanine dehydrogenase with sequences approximately > 50% similar shown.

## Discussion

In this study we compared the genomes of members of piezophilic *Colwellia* – including the most high pressure-adapted species known to date – with their piezosensitive counterparts to search for features that could confer adaptation to the deep sea. These microbes were isolated from surface and bathyal waters to abyssal and hadal depths. Both 16S rRNA gene sequence-based phylogenetic analyses and phylogenomic analyses indicate that the piezophilic *Colwellia* are closely related. While the piezophiles appear to form a single cluster based on the phylogenomic tree, in the 16S rRNA gene phylogenetic tree *C. piezophila* appears basal to not only the piezophiles but also a clade that includes piezosensitive lineages. Therefore, it is possible that piezophily has evolved multiple times within the *Colwellia.* Further whole genome sequencing will be needed to determine if all piezophilic *Colwellia* form a single clade independent from other piezosensitive microbes as has been reported for *Shewanella* (Aono *et al*., 2010). Piezophilic *Colwellia* have now been isolated from five different trenches, including the Mariana (strains MT41, MTCD1), Puerto Rico (*C. hadaliensis*), Japan (*C. piezophila*), Tonga (strain TT2012), and Kermadec (Bartlett laboratory unpublished; Lauro *et al*., 2007; Peoples *et al*., 2019a). Piezophilic members of the genus *Colwellia* are therefore widespread within deep-ocean and hadal environments.

While the piezophiles have lower coding density than their non-piezophilic counterparts, no correlation was found between genome size and optimum pressure of growth. This is in contrast to comparisons between shallow and deep pelagic datasets showing that deeper lineages appear to have larger genomes (e.g. Konstantinidis *et al*., 2009; Beszteri *et al*., 2010; Eloe *et al*., 2011; Thrash *et al*., 2014). Instead, the three piezophiles with the deepest collection depths represented some of the smallest *Colwellia* genomes examined. One possibility is that these differences reflect different selective pressures operating within seawater, sediments and amphipods. It is remarkable that strain MT41 and MTCD1, two piezophiles isolated from amphipod material in the Mariana Trench 34 years apart, share over 99% ANI. Perhaps this reflects strong selection for a particular *Colwellia* strain within the microbiome of Mariana Trench *Hirondellea gigas* amphipods, such as that seen within symbionts of deep-sea anglerfishes (Baker *et al*., 2019). Their consistent isolation from amphipods (e.g. Yayanos *et al*., 1981; Kusube *et al*., 2017) suggests that some members can be associated with hosts, and host-microbe relationships can lead to genome streamlining and smaller genome sizes (McCutcheon & Moran, 2012). Nearly all known piezophilic genera have been found in conjunction with hosts (e.g. Nakayama *et al*., 2005) and the microbial activity of the gut contents of deep-sea animals shows high levels of piezophily (Tabor *et al*., 1982). However, the genus *Colwellia* is not present in recognizable abundances within hadal amphipod metagenomes (Zhang *et al*., 2019a), their high % GC is not indicative of an endosymbiont (McCutcheon & Moran, 2012), and the obligate piezophile *Colwellia* sp. TT2012 was isolated from sediments rather than amphipods. An alternative hypothesis is that *Colwellia* may be undergoing genome reduction because of a specialized lifestyle within the deep sea, as hypothesized for some psychrophiles within sea ice (Feng *et al*., 2014). Members of this genus may instead be isolated in conjunction with amphipods because of their ability to degrade nutrient-rich decaying amphipod material, for example using genes for chitin degradation. *Colwellia* may also be ingested by amphipods as a byproduct of the feeding of these deep-sea scavenging macrofauna because of the preference of *Colwellia* for nutrient-rich particulate organic material (Hoffmann *et al*., 2017; Peoples *et al*., 2018; Boeuf *et al*., 2019).

The isoelectric point (pI) distribution of proteins within a proteome can correlate with the ecological niche of an organism (Kiraga *et al*., 2007). Here we found that piezophilic *Colwellia* have a more basic proteome than their piezosensitive counterparts. This pattern is conserved in comparisons between piezophilic and piezosensitive members of the genera *Shewanella* and *Psychromonas*, indicating it is a property that may be widespread amongst piezophiles within the *Gammaproteobacteria*. Although intracellular microorganisms also have more basic proteomes than free-living species (Kiraga *et al*., 2007), this is associated with an AT base pair enrichment not present in the piezophilic *Colwellia*. A basic proteome may be the result of the accumulation of mutations (Kiraga *et al*., 2007), consistent with the low coding density and high numbers of transposable elements within the piezophiles. Alternatively it could arise to help with charge balance within the cytoplasm, analogous to the role of the more acidic proteome of haloarchaea, which counters the high intracellular potassium ion levels present at high osmotic pressures (Paul *et al*., 2008). The intracellular inorganic and organic solute levels within piezophiles are not well known, but could be important to the maintenance of macromolecule function at high pressure (Martin *et al*., 2002; Yancey *et al*., 2001; Yancey, 2005). Among orthologous proteins piezophiles are also enriched in hydrophobic residues, including tryptophan, tyrosine, leucine, phenylalanine, histidine, and methionine. This finding has been noted in metagenomes from 4,000 m (Konstantinidis *et al*., 2009) and may be important in maintaining protein structure against water penetration at high pressure (Hummer *et al*., 1998; Somero, 2003). Specific amino acid substitutions where one amino acid is consistently replaced by another indicate that small nonpolar compounds (alanine, isoleucine), amine-containing polar compound (glutamine), and a positively charged basic compound (lysine) are selected for in piezophiles, while negatively charged acidic compounds (glutamate), polar compounds (threonine, asparagine), and non-polar compounds (valine, proline) are selected against. Similar shifts were also seen in *Desulfovibrio piezophilus* (Pradel *et al*., 2013), although different amino acids were preferentially abundant in piezothermophilic archaea (Di Giulio, 2005).

We identified a number of gene abundance characteristics that could confer adaptation to the deep ocean. Enrichments in COG J (translation), L (replication and repair), M (cell wall/membrane biogenesis), and N (cell motility) appear enriched in the piezophiles. An enrichment of category M and L has previously been observed within deep ecotypes of *Alteromonas* (Ivars-Martinez *et al*., 2008). The enrichment within the piezophiles of COG M is in part due to higher abundances of glycosyltransferases, which appear to correlate with depth within metagenome datasets (DeLong *et al*., 2006). Glycosyltransferases have been predicted to contribute to low temperature-adaptation (Methé *et al*., 2005) and could be more abundant in the psychropiezophiles because they are more stenothermic. In contrast, a fatty acid cis/trans isomerase was present only in the piezosensitive strains. The rapid cis-to-trans isomerization of unsaturated fatty acids via this isomerase has been observed in *Pseudomonas putida* P8 in response to changes in temperature and salinity (Loffeld & Keweloh, 1996; Holtwick *et al*., 1997). Furthermore, the COG category for transcription (K) is significantly enriched in non-piezophiles compared to piezophiles. This is in part due to an enrichment in the transcription factors *AraC* and *LysR*, which have a wide variety of regulatory functions including carbon metabolism and stress response (Gallegos *et al*., 1997; Maddocks & Oyston, 2008). The enrichment of COG category K in shallow-water organisms has been observed in the surface-water ecotype of *Alteromonas macleodii* (Ivars-Martinez *et al*., 2008). These findings could reflect the adaptation of non-piezophilic shallow-water microbes to a more dynamic environment, such as rapid salinity or temperature shifts associated with sea-ice or surface seawater. In contrast, autochthonous, obligate deep-ocean microbes would not be expected to experience similar rates or magnitudes of these changes.

Other specific genes biased towards piezophiles within COG M include delta-9 acyl-phospholipid desaturase and a CDP-alcohol phosphatidyltransferase. While the desaturase is upregulated at high pressure in *Photobacterium profundum* SS9 (Campanaro *et al*., 2005), this gene is present in other non-piezophilic strains of the *Colwellia* not examined here, indicating it may not be pressure-specific. An extra copy of d-alanine-d-alanine ligase is present in the piezophiles and may function in peptidoglycan biosynthesis. While this gene was reported within a flagellar operon in *Shewanella benthica* (Zhang *et al*., 2019b), in strain MT41 it is present next to a putative genomic island (Figure 5). The non-piezophile-specific copy of d-alanine-d-alanine ligase is upregulated in the proteome of strain 34H after incubation at −1°C (Nunn *et al*., 2015), perhaps reflecting a role in low temperature acclamation. Overall, the enrichment in piezophiles of genes involved in COG category M is consistent with a wealth of experimental evidence demonstrating that changes in membrane structure are critical for adapting to high hydrostatic pressure. Unsaturated fatty acids help maintain membrane structure under high pressure (Chi and Bartlett, 1995; Yano *et al*., 1998; Allen *et al*., 1999; Usui *et al*., 2012; Abe, 2013), with strain MT41 able to produce more than 15% of its total membrane fatty acids as docosahexaenoic acid (22:6; Delong & Yayanos, 1986).

Another adaptation associated with the membrane involves energetics and respiration. We identified an additional NADH ubiquinone oxidoreductase (*nuo*) gene cluster in a number of piezophiles. This unique NADH dehydrogenase, which translocates four protons per two electrons (Pinchuk *et al*., 2010), may help with energy acquisition under *in situ*, high pressure conditions. We also identified an alanine dehydrogenase specific to the piezophiles that may function in the reversible amination of pyruvate to alanine coupled with the oxidation of NADH to NAD^+^. This may act as an adaptive strategy under inhibited respiratory conditions by maintaining NADH/NAD^+^ homeostasis (Jeong & Oh, 2019), such as during shifts to anoxic conditions (Hutter & Dick, 1998; Feng *et al*., 2002) or after exposure to physical stressors impeding electron flow. Alanine dehydrogenases in *Listeria* are insensitive to inactivation up to pressures of 550 MPa (Simpson & Gilmour, 1997), transcriptionally upregulated in *Desulfovibrio piezophilus* at high pressure (Pradel *et al*., 2013), and abundant in the proteomes of strain 34H at sub-zero temperatures (Nunn *et al*., 2015). We speculate that the piezophilic alanine dehydrogenase functions in NADH/NAD^+^ homeostasis under high hydrostatic pressure conditions. In contrast, we found that TMAO reductase (*torECAD*), which reduces TMAO to TMA, was not present in any of the piezophilic *Colwellia*. A similar finding has been noted in genomes of *Psychromonas* from the guts of hadal amphipods, where the lack of TMAO reductase was attributed to the preferential need for TMAO as a piezolyte in the host amphipod over its use as an electron acceptor by the microbe (Zhang *et al*., 2018). An alternative hypothesis is that TMAO is used by microbial piezophiles as a piezolyte as it is in deep-sea metazoans (Yancey *et al*., 2001; Yancey *et al*., 2014). Finding differences in respiratory capacity within piezophiles is not unexpected. Others have previously noted the influence of collection depth and pressure on the presence and regulation of respiratory membrane-bound cytochrome c oxidases and hydrogenases (Yamada *et al*., 2000; Vezzi *et al*., 2005; Chikuma *et al*., 2007; Tamegai *et al*., 2013; Leon-Zayas *et al*., 2015; Vannier *et al*., 2015; Michoud & Jebbar, 2016; Xiong *et al*., 2016; Zhang *et al*., 2018). These changes could stem directly from pressure influences or from a greater reliance on the colonization of reduced oxygen niches associated with particles or animals (Boeuf *et al*., 2019; Peoples *et al*., 2019a). This latter possibility could be facilitated by the *tad* pilus present in the piezophilic *Colwellia* (Planet *et al*, 2003; Tomich *et al*., 2007; Pu *et al*., 2018).

Horizontal gene transfer (HGT) can provide genetic material that enhances fitness in new environments. An experimental demonstration of this impact is the introduction of a DNA photolyase gene, missing in piezophilic *Colwellia* and other deep-sea species (Delong *et al*., 2006; Lauro & Bartlett, 2008; Konstantinidis *et al*., 2009; Peoples *et al*., 2019b), into the piezophile *Photobacterium profundum* SS9 to generate a UV resistant phenotype (Lauro *et al*., 2014). It is striking that many of the *Colwellia* genes most similar to those in other piezophiles appear in clusters within variable regions that include genomic islands, putative phage genes, transposases, and toxin-antitoxin systems. Despite their smaller genome sizes, laterally transferred elements such as transposase and toxin-antitoxin genes are more abundant in the piezophilic *Colwellia* examined here, consistent with their lower coding densities. Another notable feature of these variable regions is that they differ even between closely-related strains, such as between *Colwellia marinimaniae* MT41 and *C. marinimaniae* MTCD1.

Mobile genetic elements have been suggested to confer adaptations to extreme conditions (e.g. Anderson *et al*., 2011; Pradel *et al*., 2013; Feng *et al*., 2014; Lossouarn *et al*., 2015; Mao & Grogan, 2017), such as in the known piezophile *Photobacterium profundum* SS9 (Campanaro *et al*., 2005). Deep-sea specific toxin-antitoxin systems have been identified in members of the *Shewanella* (Zhang *et al*., 2019b) and have been shown to influence the growth of *Pyrococcus yayanosii* at different pressures (Li *et al*., 2016; Li *et al*., 2018). Mobile genetic elements may provide new metabolisms within strains of *Colwellia psychrerythraea*, including the transfer of *sox* genes involved in sarcosine metabolism (Collins & Deming, 2013; Techtmann *et al*., 2016). Because of the similarity of many genomic island-associated genes in members of the piezophilic *Colwellia* to those in other gammaproteobacterial piezophiles, we suggest that HGT is a significant evolutionary process governing high pressure adaptation. Future studies should evaluate these regions and their associated genes for their importance in piezophily.

## Conclusions

In this study we compared the genomes of piezophilic and piezosensitive *Colwellia* to identify adaptations to extreme deep ocean conditions. Differences in amino acid composition, membrane and cell wall structure, respiratory capacity, tRNA modification, and complex organic carbon utilization appear to be important for life at hadal depths. Many piezophile-enriched genes are located near areas of genomic variability and could be shared among piezophiles by horizontal gene transfer. Some of the adaptations identified may not be for high pressure adaptation per se, but for lifestyles favored in hadal trenches such as affiliation with particulate organic carbon or animals.

## Materials and Methods

### Sample collection and high-pressure cultivation conditions

*Colwellia* sp. TT2012 was isolated from sediments collected via gravity core in the Tonga Trench (16° 38.505’ S, 172° 12.001’ W) at a depth of 9161 m on September 2, 2012 aboard the *R/V* Roger Revelle. Sediment from the upper three cm sediment depth horizon was mixed with filter-sterilized trench seawater and maintained at a pressure of 84 MPa and 4°C. A subset of this material was inoculated into ZoBell 2216 Marine Medium (BD Difco, Thermo Fisher, Waltham, MA, USA) under the same pressure and temperature conditions. *Colwellia* sp. TT2012 was eventually isolated as a pure culture following a number of dilution to extinction inoculations.

The isolation of both strains of *Colwellia marinimaniae* have been previously described. *Colwellia marinimaniae* MTCD1was isolated from amphipods at a depth of 10,918 m in the Challenger Deep (Kusube *et al*, 2017). *Colwellia marinimaniae* MT41 was also isolated from amphipods at a depth of 10,476 m (Yayanos *et al*., 1981). Both strains were maintained in pressurizable polyethylene transfer pipette bulbs (Samco Scientific, USA) with Zobell 2216 Marine Medium broth at 4°C and high pressure prior to sequencing.

### Pressure Sensitivity and Heavy Metal Sensitivity Testing

The growth of the strains was evaluated under different pressure and temperature conditions. Cultures of *Colwellia* strains 34H, GAB14E, and ND2E were incubated in Zobell 2216 marine medium supplemented with 100 mM HEPES and 20 mM glucose at 4°C. Growth under high hydrostatic pressure was evaluated by incubating cultures at 20 MPa increments between 0.1-80 MPa at 4°C and 16°C in triplicate. The OD600 was measured every 2.5 days for ten days. Growth rates of *Colwellia* sp. TT2012 were conducted at 0.1, 84, and 96 MPa at 4°C. Copper sensitivity tests were also performed on the piezophilic (strains MT41, MTCD1, and TT2012) and non-piezophilic *Colwellia* strains (strains 34H, GAB14E, ND2E). Copper (II) chloride dihydrate in concentrations ranging from 0 - 1.5 mM in 0.3 mM increments were added to inoculated 2216 media and the cultures were incubated at 4°C for 1-4 weeks with weekly inspection.

### Genome sequencing and assembly

Genomic DNA from *C. marinimaniae* MTCD1 was extracted from 100 mL of liquid culture after 4 weeks of incubation at 110 MPa. DNA was isolated using the Mo-Bio Ultraclean Microbial DNA Isolation Kit (Mo-Bio, USA). Genomic DNA was obtained from *Colwellia* sp. TT2012 after growth at 84 MPa and 4°C for 3 weeks. Cells were filtered onto a 0.22 um Millipore Sterivex filter cartridge (Fischer Scientific, USA) and first subjected to a lysis buffer (50mM Tris-HCl at pH 8.3, 40mM EDTA at pH 8.0, 0.75 M sucrose) and R1804M Ready-Lyse lysozyme solution (Illumina, USA). After 15 minutes of incubation at 37°C, proteinase K and sodium dodecyl sulfate were added to a final concentration of 0.5mg/ml and 1% respectively. The mixture was then incubated at 55°C for 25 minutes, followed by 70°C for 5 minutes. The lysate was treated two times with phenol-chloroform-isoamyl alcohol (24:24:1) and chloroform:isoamyl alcohol (24:1) and further purified using a Mo-Bio Utraclean DNA Isolation Kit spin column. The genomes of *C. marinimaniae* and *Colwellia* sp. TT2012 were sequenced at the Institute for Genomic Medicine (IGM) at UCSD using the MiSeq sequencing platform (Illumina, San Diego). The raw forward and reverse reads were merged using FLASH version 1.2.10 (Magoč & Salzberg, 2011) and assembled with SPAdes version 3.1.0 (Bankevich et al., 2012).

The genome of strain MT41 was sequenced to closure by whole random shotgun sequencing. Briefly, one small insert plasmid library (2–3 kb) and one medium insert plasmid library (10-15 kb) were constructed by random nebulization and cloning of genomic DNA. The sequences were assembled using the TIGR Assembler (Sutton *et al*., 1995). All sequence and physical gaps were closed by editing the ends of sequence traces, primer walking on plasmid clones, and combinatorial PCR followed by sequencing of the PCR product.

### Genomic completeness, phylogenetic analysis, and annotation

The genomes were evaluated for their completeness and phylogenetic relationships. Genome completeness and contamination was estimated using CheckM (Parks *et al*., 2015). A whole-genome phylogenetic tree was built using RAxML (Stamatakis *et al*., 2014) on the CIPRES science gateway (Miller *et al*., 2010) using the single-copy marker genes identified within CheckM. Ribosomal 16S RNA gene trees were also built by aligning sequences using the SINA Aligner (Pruesse *et al*., 2012) and built using RAxML All trees were visualized using the Interactive Tree of Life (Letunic & Bork, 2016). Genomes were annotated using the Integrated Microbial Genomes pipeline (IMG/ER; Markowitz *et al*., 2014). Pairwise average nucleotide identity between the genomes was evaluated within both the IMG interface and with orthoANI (Lee *et al*., 2016).

### Comparative genomic analysis

A comparative genomic analysis was performed between the piezophilic and non-piezophilic strains of *Colwellia* to identify whole-genome changes and specific genes unique to piezophiles. The isoelectric point (pI) of each predicted proteome was calculated using the compute pI/MW tool in the ExPASy Bioinformatics Resource Portal (Artimo *et al*., 2012). Isoelectric point values from ExPASy were rounded to the nearest tenth and the frequency of each protein pI was plotted in Figure 2a as a percent of the total proteome. Each proteome was divided into an acidic set of proteins (pI<7; N_a_) and a basic set (pI>7; N_b_) and the bias quantified using the formula ((N_b_-N_a_)/(N_b_+N_a_) × 100). The pI bias percentage is calculated such that 100% means the proteins in the entire proteome are basic, −100% means all the proteins are acidic, and 0% means equal percentage of basic and acidic proteins.

To identify specific amino acid substitutions that may correlate with piezophily, amino acid asymmetry was calculated using the procedure and software described in McDonald *et al*. (McDonald *et al*., 1999). First, proteins from the genomes were clustered using TribeMCL (Enright *et al*., 2002; scripts available at https://github.com/juanu/MicroCompGenomics) with a Blastp cutoff of 1e-5 and an inflation value of 1.4. Orthologous single-copy gene clusters present in both the piezophiles and *Colwellia psychrerythraea* 34H were aligned using MAFFT (Katoh & Toh, 2008) and then processed with the Asymmetry programs AmbiguityRemover (using a value of 2 for the number of adjacent sites), AsymmetryCounter, and AsymmetryScaler (with three decimal places and 100 replicates; McDonald *et al*., 1999). Approximately 346,000 aligned amino acid sites were examined in each comparison. Comparisons were also performed between the *Shewanella* strains *S. benthica* KT99, *S. violacea* DSS12, and *S. piezotolerans* WP3 against the piezosensitive *S. sediminis* EB3.

Protein abundances from the genomes were compared to identify attributes preferentially enriched in either the piezophiles or piezosensitive strains. General COG category distributions were evaluated using IMG/ER annotations. For the identification of differentially-abundant specific proteins, protein clusters were generated using the TribeMCL analysis as described above. These identified protein clusters were further screened using blastp (Altschul *et al*., 1990) against the nr database for their prevalence in other *Colwellia* genomes, other piezophile genomes, or other metagenomes. This manual curation allowed for the identification of both genes differentially abundant within the groups of genomes immediately discussed here but also allowed for a culled, smaller dataset of genes that may be present in other deep-ocean isolates and datasets.

Certain genomic features within the genomes were also identified. Genomic islands were identified using IslandViewer (Bertelli *et al*., 2017). Regions that may represent genomic islands were also identified using the Mean Shift Genomic Island Predictor (MSGIP; de Brito *et al*., 2016). As incomplete genomes appeared to give spurious results, the total number of genomic islands are reported only for the complete genomes of *Colwellia marinimaniae* MT41 and *C. psychrerythraea* 34H. However, genomic islands for some of the partial genomes are shown here (e.g. Figure 4, Figure 5) only when IslandViewer or MSGIP identified a region as a genomic island, it was in a similar region as a genomic island found in either of the 34H or MT41 genomes, and it appeared to be a region of variability based on IMG/ER annotations. The homology of these variable regions was analyzed using blastn and visualized with the R package genoPlotR (Guy *et al*., 2010) and Kablammo (Wintersinger *et al*., 2015). Putative transposases and toxin/antitoxin genes were identified based on IMG/ER annotations. Putative viral regions of each genome were also identified based predominantly on IMG/ER annotations with a functional search using the terms ‘phage’ and ‘virus,’ but also with VirFinder (Ren *et al*., 2017) and VirSorter (Roux *et al*., 2015). Different types of flagella and pili were annotated using MacSyFinder and TXSScan (Abby *et al*., 2016; https://galaxy.pasteur.fr/#forms::txsscan) with default parameters. Carbohydrate-active enzymes within each genome were identified using dbCAN (Yin *et al*., 2012).

## Declarations

### Ethics approval and consent to participate

Not applicable.

### Consent for publication

Not applicable.

### Availability of data and material

The genome sequences of strains MT41, MTCD1, and TT2012 have been deposited at GenBank under the accessions CP013145, GCA_001432325, and GCA_001440345, respectively. The assembled and annotated genomes of strains MT41, MTCD1, and TT2012 can be located in IMG/JGI under the IMG taxon IDs 2501651205, 2585427605, and 2585428047 respectively.

## Authors’ contributions

LMP, TSK, JAU, KM, RAC, AAY, BAM performed experimental and bioinformatics work. LMP, TSK, DHB performed data analysis. LMP, TSK, DHB wrote the manuscript. All authors read and approved the final manuscript.

## Acknowledgements

We thank Stephen Techtmann and Terry Hazen for providing the *Colwellia psychrerythraea* strains GAB14E and ND2E. Thanks to Priya Narasingarao and Sheila Podell for constructive input. We appreciate the support of the crew of the *R/V Revelle* and those associated with the MOTT expedition for their help collecting these samples. This includes but is not limited to Rosa Leon-Zayas, Jenan Kharbush, Rachael Hazael, and Fabrizia Foglia.

