## Supplementary Figures for "Distinctive Gene and Protein Characteristics of Extremely Piezophilic *Colwellia*"

Supplementary Information, bioRxiv

Supplementary Figure 1. Growth curves of strains of *Colwellia psychrerythraea* as a function of pressure and temperature. A, *C. psychrerythraea* 34H; B, *C. psychrerythraea* ND2E; *C. psychrerythraea* GAB14E.

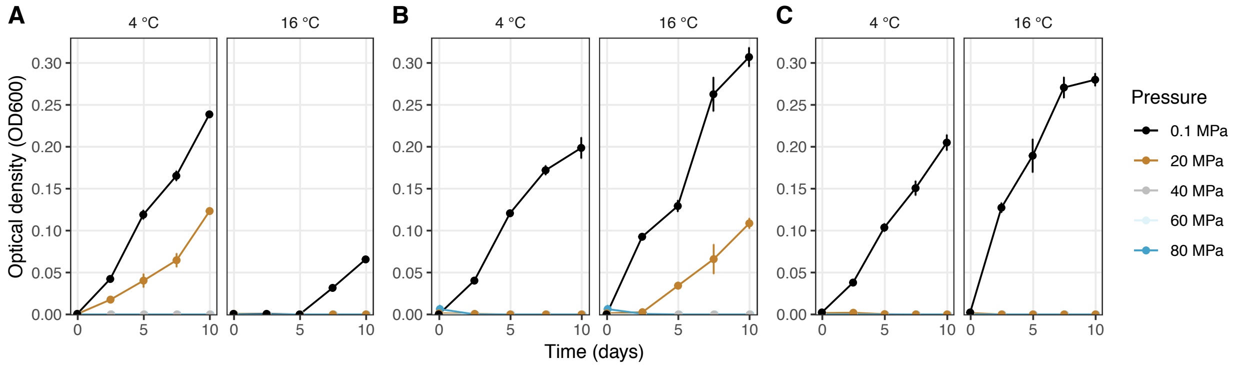

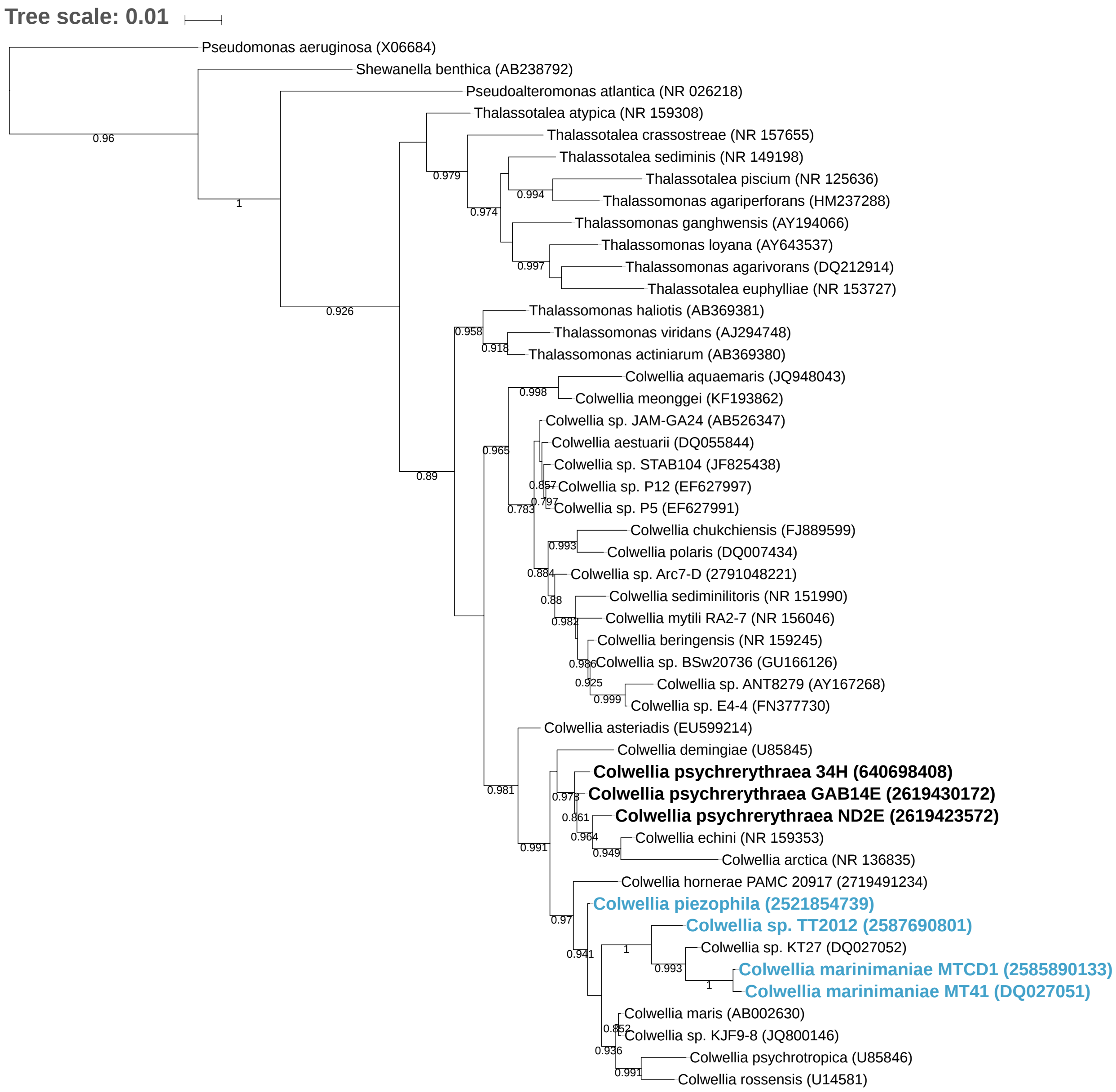


Supplementary Figure 2. Ribosomal 16S RNA gene tree of members of the *Colwellia*. Strains in bold were compared in this study; blue are piezophilic, black are piezosensitive.

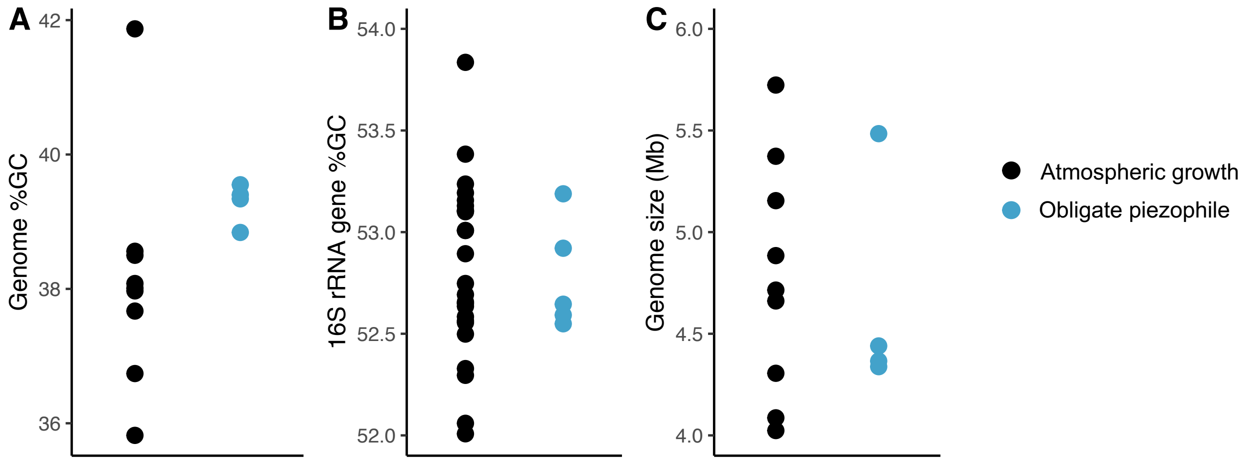


Supplementary Figure 3. Strains of *Colwellia* as a function of (A) their genome % GC, (B) full length 16S rRNA gene % GC, and (C) genome size.

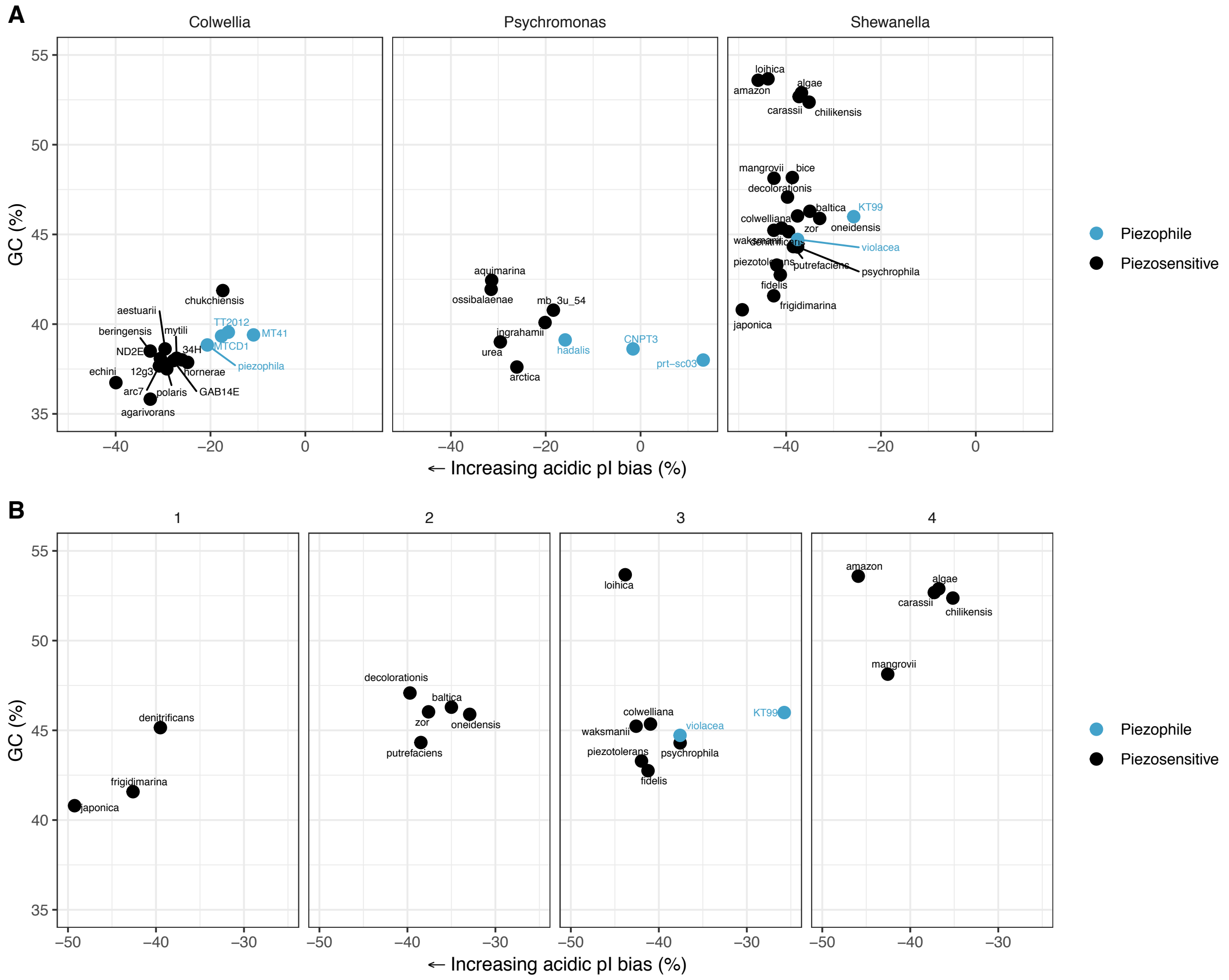


Supplementary Figure 4. Isoelectric point (pI) bias of proteins within each strain. A; Isoelectric point (pI) bias of proteins plotted as a function of % GC content within the genomes of members of the *Colwellia*, *Psychromonas*, and *Shewanella*. B; pI bias of members of the *Shewanella* plotted as a function of % GC when taking into account within-genus position broadly based on a phylogenetic tree generated by Alex & Antunes, 2019. Strains are organized by general clade grouping, labeled here as clades 1-4.

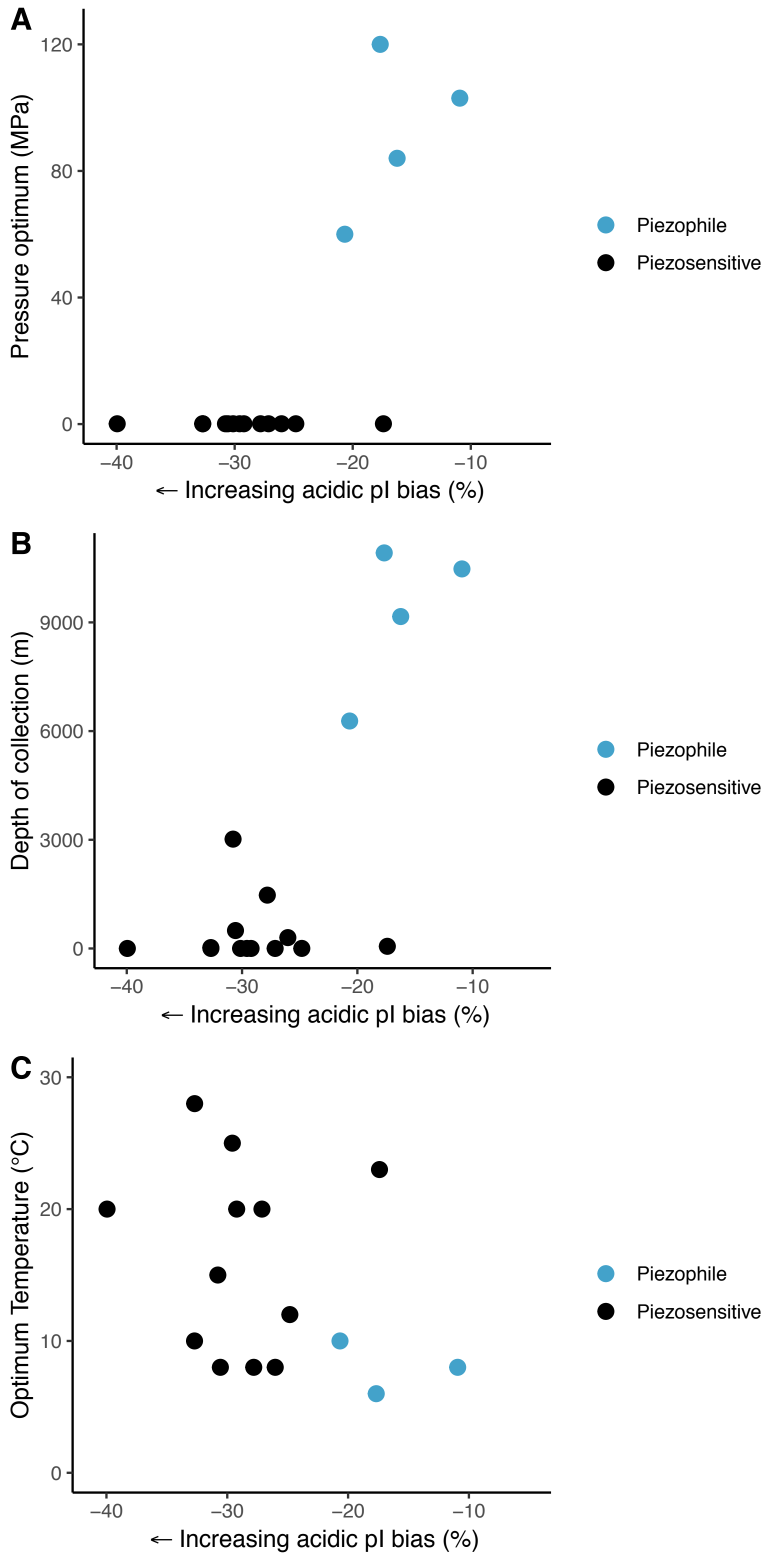


Supplementary Figure 5. Isoelectric point bias of members of the *Colwellia* as a function of optimum pressure of growth (A), depth of collection (B), or optimum temperature (C).

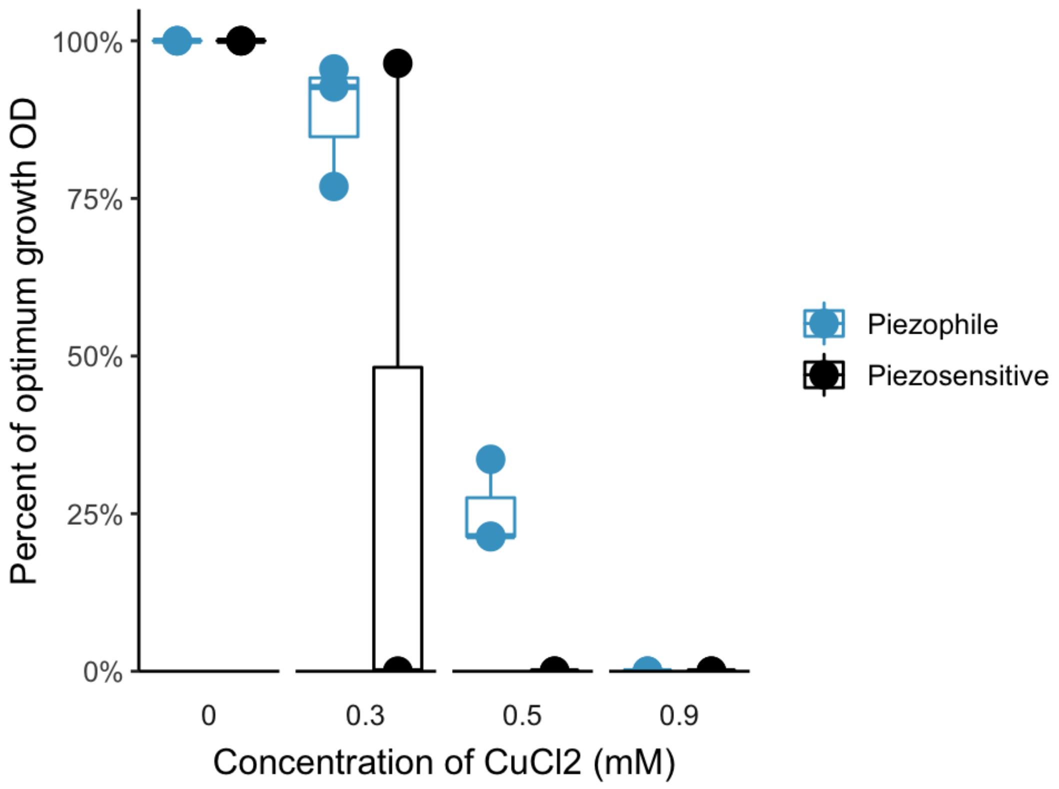


Supplementary Figure 6. Growth of strains of *Colwellia* (piezophile, n=3; piezosensitive, n-=3) measured as optical density (OD600) as a function of copper (II) chloride dihydrate concentrations at 4°C and optimum pressure. The OD values are plotted as a percentage of the OD measured under optimal conditions, when no copper (II) chloride is added.

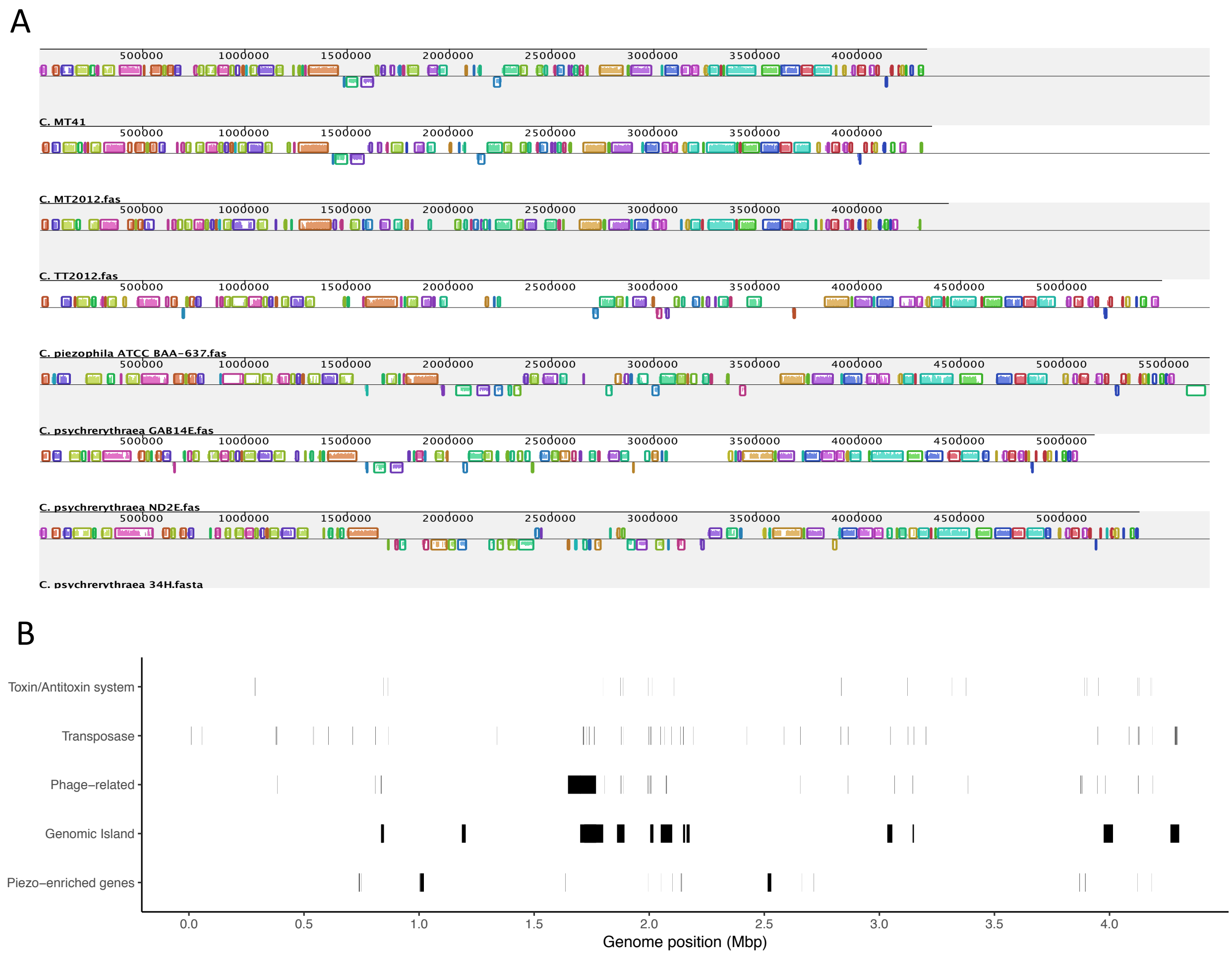


Supplementary Figure 7. A; Mauve alignment of strains of *Colwellia*. *C. marinimaniae* MTCD1 is labeled as ‘C. MT2012’. B; Locations of specific genomic elements identified within *Colwellia marinimaniae* MT41.
